# Semaphorin 3A binding to chondroitin sulfate E enhances the biological activity of the protein, and cross-links and rigidifies glycosaminoglycan matrices

**DOI:** 10.1101/851121

**Authors:** Lynda Djerbal, Romain R. Vivès, Chrystel Lopin-Bon, Ralf P. Richter, Jessica C.F. Kwok, Hugues Lortat-Jacob

## Abstract

Semaphorin 3A (Sema3A) is a secreted protein that signals to cells through binding to neuropilin and plexin receptors and provides neurons with guidance cues key for axon pathfinding, and also controls cell migration in several other biological systems. Sema3A interacts with glycosaminoglycans (GAGs), an interaction that could localize the protein within tissues and involves the C-terminal domain of the protein. This domain comprises several furin cleavage sites that are processed during secretion and in previous works have hampered recombinant production of full-length wild type Sema3A, and the biochemical analysis of Sema3A interaction with GAGs. In this work, we have developed a strategy to purify the full-length protein in high yield and identified two sequences in the C-terminal domain, KRDRKQRRQR and KKGRNRR, which confer to the protein sub nM affinity for chondroitin sulfate and heparan sulfate polysaccharides. Using chemically defined oligosaccharides and solid phase binding assays, we report that Sema3A recognizes a (GlcA-GalNAc4S6S)_2_ motif but not a (GlcA2S-GalNAc6S)_2_ motif and is thus highly specific for type E chondroitin sulfate. Functionally, we found that Sema3A rigidified CS-E films that mimic the GAG presentation within extracellular matrices (ECMs), suggesting that Sema3A may have a previously unidentified function to cross-link and thus stabilize GAG-rich ECMs. Finally, we demonstrated that the full-length Sema3A is more potent at inhibiting neurite outgrowth than the truncated or mutant forms that were previously purified and that the GAG binding sites are required to achieve full activity. The results suggest that Sema3A can rigidify and cross-link GAG matrices, implicating Sema3A could function as an extracellular matrix organizer in addition to binding to and signaling through its cognate cell surface receptors.

## INTRODUCTION

Semaphorins form a large family of either secreted or membrane associated signaling proteins initially described as repulsive axon guidance clues, but now known to control many other biological processes, such as angiogenesis, bone turnover and homeostasis, cardiac development, immune response and associated disorders, and cancer [1, 2]. Within vertebrates, about 20 semaphorins have been identified and classified into five groups (semaphorins 3 to 7) [3]. They all consist of a 500 amino acid long N-terminal Sema domain, providing a non-covalent dimerization interface, followed by a plexin-semaphorin-integrin (PSI) domain, an Ig-like domain (excepted classes 5 and 6) and class-specific additional sequences in the C-terminus [4-6].

Semaphorins, most of which are transmembrane or glycophosphatidylinositol membrane anchored proteins, interact with plexin receptors exposed on adjacent cells to mediate short-range/paracrine signals, a mechanism based on receptor di-or oligo-merization and downstream activation of the intracytoplasmic GTPase-activating protein domain [6-8]. In contrast to all other vertebrate semaphorins, class 3 semaphorins (which comprises 7 members, A to G), are secreted molecules that can diffuse in the extracellular environment, potentially mediating more long range signaling activities. In addition, the binding of class 3 semaphorins to plexin is facilitated by neuropilin (Nrp) that interacts with both proteins, thereby strengthening the semaphorin-plexin complex [9].

Semaphorin 3A (Sema3A), the best characterized member of the secreted vertebrate semaphorins, also binds to glycosaminoglycans (GAGs). This interaction could contribute to localize Sema3A appropriately within tissues, for example, in establishing chemorepulsive boundaries in specific areas [10]. At the cellular level, it has been observed that neurons expressing GFP tagged Sema3A displayed a punctuate distribution of the protein at the surface of both axons and dendrites, which could be eliminated following treatment with an excess of GAGs or with chondroitinase ABC [11], suggesting an interaction of Sema3A with chondroitin sulfate (CS). In the adult central nervous system (CNS), Sema3A also co-localizes with CS [10, 12, 13] such as within perineuronal nets (PNNs) [12], dense extracellular matrices that enwrap the soma and dendrites of specific neurons [14].

PNNs stabilize synaptic connections during development, closing the critical period of brain plasticity during which experience can shape neural circuitry. Preventing PNN formation increases axonal sprouting, synaptic plasticity and memory retention in adult mice, which, interestingly, can also be observed following localized digestion with chondroitinase ABC [15, 16]. Enzymatic removal of CS from the PNNs extends the period of plasticity in models of visual cortex plasticity [17] and it was recently shown that Sema3A accumulation correlates with PNN maturation, that is, Sema3A appears in the PNN at the closure of the critical period [18, 19]. Enzymatic removal of CS also promotes axon regeneration following spinal cord injury [20]. The role of Sema3A in these processes, and its association to CS, are not fully clear, but its presence within the extracellular matrix (ECM) could be an important contributing factor to the stabilization of synaptic contacts in the CNS but also to the failure of axonal regrowth beyond the glial scar in the injured spinal cord.

The present work was designed to better characterize the biochemical and functional features of the Sema3A/CS interaction. A number of studies have already suggested that Sema3A binding to GAGs is mediated by the C-terminal domain of Sema3A [11, 21]. This domain features two internal furin-like proprotein convertase cleavage sites (R-X-[K/R]-R) at positions 552-555 and 758-761 (site 1 and 2; Figure 1), and their processing either enhances (site 2) or suppresses (site 1) the chemorepulsive activity of the protein [22]. The full-length Sema3A monomer, which has a molecular mass of 90 kDa, is also processed upon overexpression, with cleavage at site 1 giving rise to a 65 kDa protein comprising the entire Sema domain plus a large part of the PSI domain but lacking a small part of the PSI domain, the Ig-like domain and an unstructured C-terminal basic tail which contains a cysteine at position 723 for covalent disulfide bonding of the Sema3A dimer [23]. A strategy that has been commonly used to prevent this processing and to produce full length Sema3A is by introducing mutations at the furin sites [5, 24, 25]. Mutation at site 1 produces a full length Sema3A bearing a strong chemorepulsive activity in neuronal outgrowth assay. It is, however, unclear if furin site 2 was cleaved and involved in the activity. Due to the difficulties in the purification of full-length wild type (WT) Sema3A, the importance of the C-terminal domain in Sema3A for GAG binding and bioactivity has not yet been studied. To address these aspects, we have designed here a specific protocol to successfully purify in high yield the full-length WT Sema3A overexpressed in HEK293-EBNA cells [26]. Using this purified WT Sema3A, we identified the GAG binding sites and characterized their involvement in the biological activity of the protein. Conversely, we also investigated the features of GAG structures required for Sema3A binding using solid phase binding assays by surface plasmon resonance (SPR) and quartz crystal microbalance (QCM-D). Results indicate that Sema3A binding to CS is sulfation pattern-dependent, with binding strengths in the pM range for CS-E polysaccharides. With the use of synthetic CS oligomers, we discovered that the minimal structure that Sema3A recognized with high affinity and specificity is a CS-E tetrasaccharide. Competition assays further indicate that the strong binding of Sema3A arises from multivalent interactions within CS-E rich matrices, which provides a putative mechanism for the selective binding of Sema3A to PNNs. Our data also indicate that Sema3A can rigidify and cross-link GAG matrices, implicating Sema3A could function as an extracellular matrix organizer in addition to binding to and signaling through its cognate cell surface receptors.

**Figure 1:**
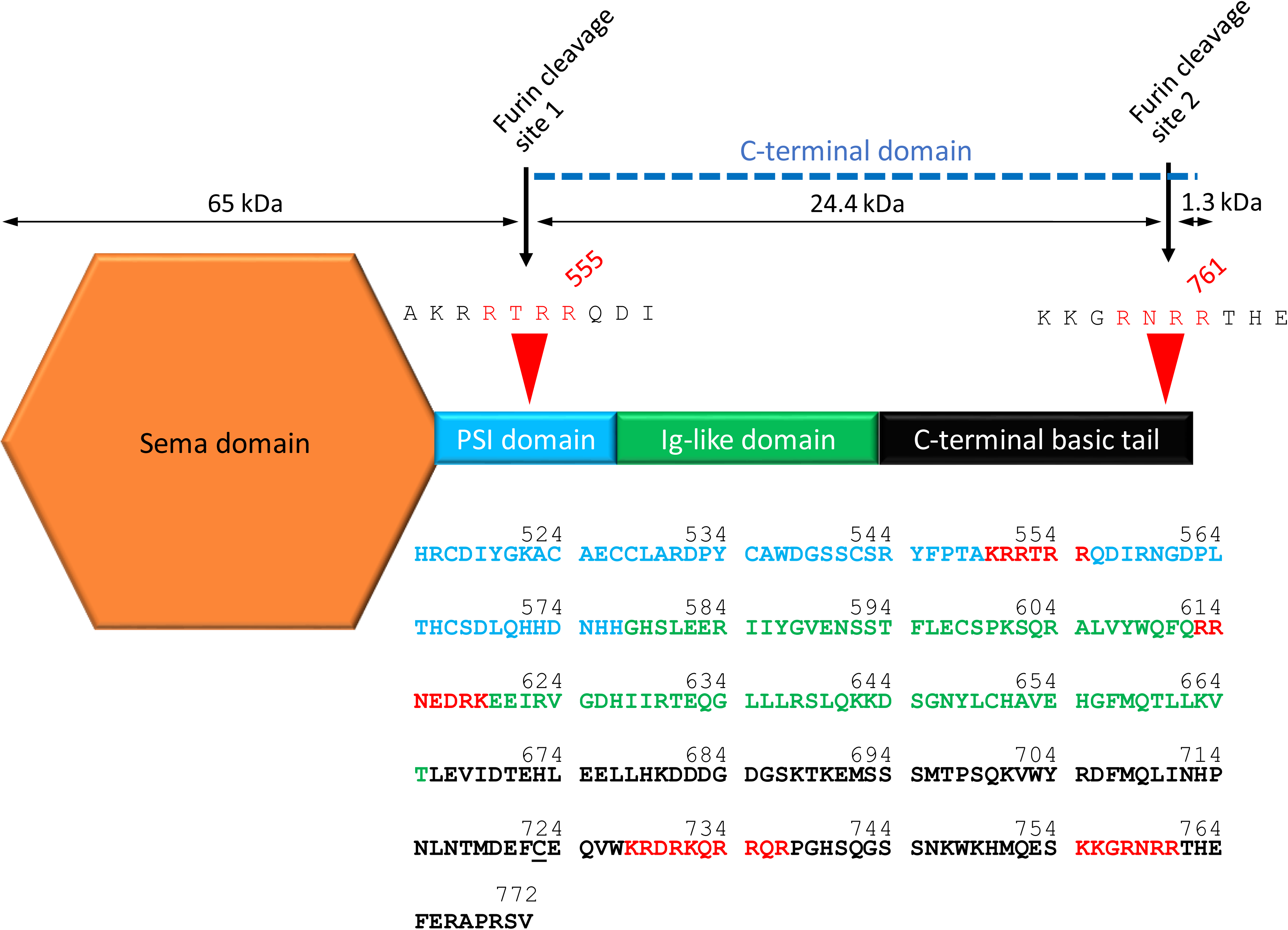
Mapping of the Sema3A heparin binding domain. Schematic representation of Sema3A, highlighting the Sema, PSI and Ig-like domains (in orange, blue and green respectively) followed by the unstructured C-terminal basic tail (in pink). The sequence of the protein, downstream the Sema domain, is indicated with the same color code and shows the four clusters of basic amino acids. The two furin cleavage sites, and the C-terminal domain which encompasses the amino acids part after the first cleavage site, are also indicated. The cysteine that forms a disulfide bond in the Sema3A homo-dimer is underlined.

## MATERIALS AND MATHODS

### Expression and purification of Sema3A wild type and mutant proteins

Rat Sema3A (residues 23-772, missing its own N-terminal signal peptide and the two first amino acids of the Sema domain) was cloned downstream of the CMV promoter into the pTT22SSP4 expression vector, comprising the Secreted Alkaline Phosphatase signal peptide and a 8× His tag. Mutants of Sema3A were generated from pTT22SSP4-Sema3A by PCR quick change. Wt and mutant proteins were expressed from HEK-6E EBNA cells growing in suspension (NRCC, Biotechnology Research Institute). For that purpose, cells were transfected with 250 µg of Sema3A vector using 500 µl of 1 µg/µl linear polyethylenimine (PEI, Polysciences, 23966-1) solution. HEK-EBNA-Sema3A cells (1.7 × 10^6^) were grown for 4 days in 250 ml of FreeStyle F17 Expression Medium supplemented with 4 mM L-glutamine, 0.1 % kolliphor P188 (Sigma, K4894) and 25 µg/ml G418 in a humidified incubator at 37°C with 5% CO_2_. The medium was supplemented with 0.5 % Tryptone TN1 (Organotechnie, 19553) at 24 h post-transfection and one tablet of EDTA-free protease inhibitor cocktail (Roche, 5056489001) was added at 1 and 3 days post-transfection. To purify the Sema3A fraction that remained associated to the cell surface, the cell pellet was washed in 10 mM phosphate-1 M NaCl pH 7.2 and the supernatant loaded onto a 1 ml Ni-NTA chromatography column. We found that some of the mutants of Sema3A, and the truncated Sema3A, were secreted into the medium. To purify these proteins, medium was collected, dialyzed 3 times against 2 L of PBS (3.5 kDa cutoff) and injected on the Ni-NTA column. The column was washed with 50 ml of PBS-1 M NaCl, 50 ml of PBS-0.5 % triton and then PBS-20 mM imidazole. The protein was eluted with 10 ml of PBS-300 mM imidazole. The material was next purified by cation-exchange chromatography on a 1 ml sulfo-propyl-sepharose column equilibrated in PBS and eluted with a 0.15-1 M NaCl gradient.

### Identification of GAG binding sites

Sema3A GAG binding sites were mapped essentially as previously described [27]). Briefly, heparin beads were activated with 200 mM of l-ethyl-3-(3-dimethylaminopropyl) carbodiimide (EDC) and 50 mM of N-hydroxysuccinimide (NHS), in 50 mM MES, 150 mM NaCl pH 5.5, for 10 min at room temperature under stirring. Excess EDC/NHS reagents were removed by 3 steps of centrifugation/PBS wash. Heparin beads were next incubated with Sema3A (from 20 to 180 µg) for 2 h at room temperature, under gentle agitation, and the reaction was then stopped by incubation for 10 min in 1 M Tris, pH 7.5 (100 mM Tris final concentration). After washing the beads twice with 2 M NaCl PBS, covalently bound Sema3A was denatured with 2 M urea and 75 mM β-mercaptoethanol, at 100 °C for 1 h, prior to overnight digestion with thermolysine (50 mIU) at 50 °C. Released peptides were removed by washing the beads 3 times with PBS supplemented with 2 M NaCl, 929.5 mM β-mercaptoethanol and 1% triton, while cross-linked peptides, overlapping the Sema3A GAG binding sites, were identified by Edman degradation automated sequencing performed directly on the beads.

### GAG disaccharide analysis

GAG disaccharide analysis was performed as previously described [28]. CS in 50 mM Tris-HCl pH 7.5, 50 mM NaCl, 2 mM CaCl_2_ was digested with 500 mU of chondroitinase ABC for 24 h at 37°C. Composition was determined by RPIP-HPLC, using a Phenomenex Luna 5 µm C18 reversed phase column (4.6 × 300 mm; Phenomenex, Le Pecq, France) heated at 54°C and equilibrated at 1.1 mL/min in 1.2 mM tetra-N-butylammonium hydrogen sulfate (TBA) and 8.5% acetonitrile, to which was applied a multi-step NaCl gradient (0–30 mM in 1 min, 30–90 mM in 39 min, 90–228 mM in 2 min, 228 mM for 4 min, 228-300 mM in 2 min, 300 mM for 4 min) calibrated with CS disaccharide standards. On-line post-column disaccharide derivatization was achieved by the addition of 2-cyanoacetamide (0.25%) in NaOH (0.5%) at a flow rate of 0.35 mL/min, followed by fluorescence detection (excitation 346 nm, emission 410 nm).

### GAG biotinylation

CS-D purified from shark cartilage and with an average molecular mass of 38 kDa was obtained from Seikagaku (400676). CS-E purified from squid cartilage (30 kDa) was obtained from Amsbio (AMS.CSR-NACS-E2.SQC-10). HS purified from porcine intestinal mucosa with an average molecular mass of 12 kDa and an average sulfation of 1.6 per disaccharide was obtained from Celsus Laboratories (Cincinnati, OH, USA; Supplementary Fig.3). These GAGs were biotinylated at their reducing end, essentially as described previously [29, 30]. Briefly, CS-D, CS-E and HS at 0.2-0.5 mM were reacted for 24 h at 37°C with 10 mM of EZ-link alkoxyamine-PEG4-SS-PEG4-biotin (ThermoScientific, 26138), in 50 mM acetate buffer pH 4.8 and 20 mM aniline, after which the mixtures were extensively dialyzed against water to remove unreacted biotin, and freeze dried. Biotinylation was controlled by dot blotting the material onto a Nylon transfer membrane (Amersham Life Sciences, Hybond-N+), which was revealed using extravidin-peroxidase.

### Preparation of synthetic, size-defined CS-D and CS-E oligosaccharides

CS-D and CS-E oligosaccharides were obtained from a single disaccharide precursor bearing a benzylidene acetal on the GalN moiety and a naphylmethyl group as aglycon that was prepared by semisynthesis from commercially available chondroitin sulfate polymer as previously described [31]. Di-, tetra-and hexa-saccharides were then synthesized after several selective protection and deprotection steps followed by sulfation of hydroxyl group and deprotection as previously described [29]. For some assays, CS-E oligosaccharides were biotinylated. For that purpose, a 2-benzyloxycarbonylaminoethyl group was introduced on the key disaccharide and after similar steps of selective protection/deprotection, sulfation, biotinylation and deprotection, the biotinylated oligosaccharides were isolated as described [33]. The resulting molecular masses for the CS-E di-, tetra-and hexa-saccharides were 1006 Da, 1611 Da and 2217 Da, respectively.

### Surface plasmon resonance (SPR) based binding assays

Biacore 3000 and Biacore T200 instruments (both GE Healthcare) were used for SPR analysis of Sema3A binding to immobilized GAGs, and for competition experiments, respectively. Four flow cells of a CM4 sensor chip (GE Healthcare) coated with a dextran film were activated with 50 µL of 400 mM N-ethyl-N’-(diethylaminopropyl)-carbodiimide (EDC) and 50 µL of 100 mM N-hydroxysuccinimide (NHS) after which 50 µl of streptavidin (Sigma, S0677; 60 kDa) at 80 µg/ml in 10 mM sodium acetate buffer, pH 4.2 was injected. Remaining activated groups were blocked with 50 µl of 1 M ethanolamine, pH 8.5. This procedure allowed coupling ~1000 resonance units (RU) of streptavidin on each flow cell.

Biotinylated CS-D, CS-E and HS polysaccharides (5 µg/ml in 10 mM Hepes, 0.3 M NaCl, pH 7.4) were captured to a level of ~60 RU, each on one surface, and the fourth surface was left untreated and served as negative control [29]. This sensor chip functionalization strategy effectively generates films that reproduce salient features of GAG-rich cell coats: anchorage of the GAGs via their reducing end to streptavidin replicates the topology of proteoglycans, with one or several GAGs attached to a core protein; moreover, the dextran film into which the GAG-streptavidin complexes are embedded represents a hydrated GAG-rich matrix such as found in PNNs and other cell coats.

A sensorchip comprising size-defined synthetic CS-E di-, tetra-and hexa-saccharides was made similarly, also with a capture level of ~60 RU. From the molecular masses and capture levels, we can estimate that each streptavidin displays between 2 and 4 oligosaccharides on average depending on oligosaccharide size.

Before use, the chip was submitted to several injections of HBS buffer containing 2 M NaCl, and was then washed by continuous flow of HBS-P buffer (Hepes 10 mM, NaCl 0.15 M, P20 detergent 0.05%, pH 7.4). For binding assays, a range of concentrations of wild type or mutant Sema3A (250 µl) was injected at 25 °C and 60 µl/min over both negative control and GAG-presenting surfaces after which the surfaces were regenerated with a 250 µL pulse of 2 M NaCl. Control sensorgrams were subtracted online from GAG sensorgrams. The data were analyzed by fitting both the association and dissociation phases for each Sema3A concentration to a 1:1 binding model including a mass transfer step using the BIAevaluation 3.1 software (GE Healthcare). Kinetic constants represent the mean and standard errors based on two independent experiments per condition. For competition studies, Sema3A was preincubated with a range of concentrations of either CS-D or CS-E tetrasaccharides and injected over the CS-E tetrasaccharide functionalized surface.

### Quartz crystal microbalance with dissipation monitoring (QCM-D) measurements

A Q-Sense E4 system (Biolin Scientific, Västra Frölunda, Sweden) was used for QCM-D analysis of Sema3A binding to GAG films as previously described [32]. Briefly, a dense monolayer of streptavidin was first formed on a supported lipid bilayer (formed by spreading sonicated unilamellar vesicles made from a 95:5 (mol:mol) mixture of dioleoylphosphatidylcholine (DOPC) and dioleoylphosphatidylethanolamine-CAP-biotin in 10 mM Hepes, 150 mM NaCl, pH 7.4) on four silica-coated QCM-D sensors (QSX303; Biolin Scientific). Biotinylated GAGs were then captured on 3 of these sensors (the fourth one being a negative control) and the buffer was exchanged with 10 mM phosphate-220 mM NaCl, pH7.2. This sensor functionalization strategy generates a film of GAGs that are anchored *via* their reducing ends, akin to their anchorage in proteoglycans, and at saturation, the mean distance between GAG chains is approximately 7 nm [32], comparable to the size of Sema3A. Wild type Sema3A proteins: Sema3A-90 (20 µg/ml in PBS containing ~290 mM NaCl) and Sema3A-65 (20 µg/ml in PBS containing ~200 mM NaCl) were injected over the assembled film for 20 min at 20 µl/min, after which they were eluted with 4 M of guanidine hydrochloride. Shifts in frequency, Δ*f*, and dissipation, Δ*D*, were recorded at six overtones (*n =* 3, 5, 7, 9, 11 and 13). Data for the normalized frequency shift, Δ*f* = Δ*f*^n^/ *n*, and Δ*D* at the 5^th^ overtone (*n* = 5) are presented here; all other overtones showed qualitatively equivalent responses.

### Functional assay

Three month old female Sprague Dawley rats were sacrificed. Dorsal root ganglion (DRG) neurons were extracted from the spinal column of each animal and dissociated into individual neurons with 0.2% collagenase (Sigma, C9407) and 0.1% trypsin (Sigma, T0303) for 10 min at 37°C. These isolated cells were subjected to density gradient centrifugation (1000 × *g* for 15 min) with DMEM-15 % BSA. Pelleted alive cells were resuspended in 2 ml of DMEM-1 % insulin-transferrin-selenium (ITS, VWR, 734-1315) - 1% penicillin-streptomycin-fungizon (PSF, Invitrogen, 15240062) - 0.1% NGF (Sigma, N2513) and centrifuged at 2000 × *g* for 2 min. DRG cells were distributed over a 24 well plate (500 µl/well) containing glass coverslips coated beforehand with 1 ml/well of 20 µg/ml poly-D-lysine (Sigma, P1149) overnight and then with 500 µl/well of 1 µg/ml laminin (Sigma, L2020) for 1 h. DRG neurons were cultivated in a humidified incubator at 37°C with 5% of CO_2_ for 24 h. Wild type Sema3A was added at final concentrations of 0.125, 0.5 and 2 µg/ml, while C1-, C2-, C3- or C4-Sema3A were added at a final concentration of 2 µg/ml. After 48h, the cells were immuno-stained with an antibody against β-tubulin III (Sigma, T8660; 1:500). The slides were examined with an epifluorescence microscope (V-M4D, Olympus and Perkin-Elmer). The images were recorded with 20× magnification and with a sCMOS camera (Hamamatsu Orca Flash4, Volocity). The number of neurons with neurites (defined as projections with a length of at least twice the soma diameter) were quantified. Data were obtained from culture of 3 rats, 3 coverslips per rat and 5 views per coverslip. The percentage of neurons with neurites were counted. A one-way Anova test was used to confirm the variation between different conditions.

## RESULTS

### Semaphorin 3A expression and purification

Sema3A comprises two internal furin-like proprotein convertase cleavage sites (R-X-[K/R]-R) at positions 552-555 and 758-761 (Fig. 1). It is usually processed during overexpression giving rise to a protein comprising the entire Sema domain plus a large part of the PSI domain but lacking the Ig-like domain and the C-terminal tail, unless mutations are introduced to prevent this processing [9]. Initial sequence analysis however suggested that potential CS-binding sites are located in the C-terminal domain. Here, to produce full length and wild-type Sema3A, we designed a specific protocol in which N-terminally His-tagged Sema3A was expressed in high yield from HEK293-EBNA cells transfected with a vector containing viral sequences allowing for episomal DNA replication [26]. Conditioned medium was collected after 4 days (i.e. the time at which immunoblotting with an antibody directed against the His-tag showed maximal intensity, data not shown) and the Sema3A was purified using Ni-NTA affinity chromatography. SDS Page electrophoresis of the resulting material (Sup mat Fig. 1A) showed a major band of ~ 65 kDa (referred to as Sema3A-65) and a minor ~ 90 kDa form (referred as Sema3A-90). The existence of the 65 kDa form, presumably corresponding to a furin processed Sema3A was already reported in both embryonic mouse [22] and adult rat brain [12]. This material was not retained by cation exchange chromatography onto a sulfo-propyl-sepharose column and was purified to homogeneity using gel filtration chromatography. Interestingly, treating the overexpressing cells with high salt containing buffer (PBS supplemented with 1 M NaCl), released a large amount of the 90 kDa protein, which could be purified using Ni-NTA chromatography and SP-sepharose column, from which it was eluted with ~ 700 mM NaCl (Sup mat Fig. 1B). Altogether, this suggests that, upon expression, the majority of the full-length Sema3A is retained at the cell surface by its C-terminal domain, presumably through interaction with negatively charged GAGs [12], while a fraction of the expressed protein is processed and released as a 65 kDa soluble form in the extracellular medium.

### Identification of the GAG binding sites

As electrostatic forces usually contribute significantly to protein-GAG interactions, binding sites are commonly enriched in basic residues. Examination of the C-terminal domain of Sema3A revealed the presence of four such basic sequences, herein referred to as clusters 1 to 4, two of which (clusters 1 and 4) being also furin cleavage sites. These clusters, KRRTRR (residues 550-555), RRNEDRK (residues 613-619), KRDRKQRRQR (residues 728-737) and KKGRNRR (residues 755-761) are distributed across the PSI domain, the Ig like domain, and the downstream unstructured C-terminal tail of the protein (Fig. 1).

To investigate the possible involvement of these clusters in GAG recognition, we first used a mapping strategy based on the formation of cross-linked heparin-protein complexes, the proteolytic digestion of these complexes and the identification of the polysaccharide-covalently bound peptides by N-terminal sequencing [27]. Noteworthy, amino acids involved in cross-linking with the polysaccharide could be easily identified, as saccharide conjugation prevented their elution, which resulted in a “gap” within the detected sequences. Analysis was performed on Sema3A-90, and on Sema3A-65 which served as negative control to rule out possible irrelevant detected peptides (data not shown). Results for Sema3A-90 yielded the sequences FQRRNEDRKEE, VW**K**RDR and MQES**KK**G encompassing clusters 2, 3 and 4, respectively (Fig. 1). However, none of the cluster 2 containing sequences exhibited the expected missing K residue that acknowledged actual protein/polysaccharide cross-linking, as indicated in bold for clusters 3 and 4. These results therefore highlighted clusters 3 and 4 as putative heparin binding domains, suggested no involvement of cluster 1 and provided inconclusive data on cluster 2.

To further analyze the respective contribution of these clusters in GAG binding, we designed mutants in which [KRDRKQRRQR] and [KKGRNRR] were deleted (C3-and C4-Sema3A, respectively). Attempts to eliminate clusters 1 and 2 in the Sema3A sequence yielded a protein that remained intracellular (data not shown). Point mutations were thus introduced, including [KRRTRR] to [ASSTSA] mutations (C1-Sema3A) and [RRNEDRK] to [SANEDRK] mutations (C2-Sema3A) to disrupt the GAG recognition consensus sequences. C1-and C2-Sema3A expression and purification displayed the same features as WT-Sema3A, i.e, they remained associated to the cell surface from which they could be released using high salt washes, and they eluted from cation exchange chromatography with 700 mM NaCl. The only observed difference was the absence of the Sema3A-65 form in C1-Sema expression, consistent with cluster 1 being a furin cleavage site. In contrast, C3-and C4-Sema were released in soluble forms in the supernatant of expressing cells and only required ~ 450 mM of NaCl for their cation exchange chromatography elution (Sup mat Fig. 2). These data, together with those of the mapping approach, thus suggest a major involvement of clusters 3 and 4 in GAG binding and cell surface/matrix retention of Sema3A.

**Figure 2:**
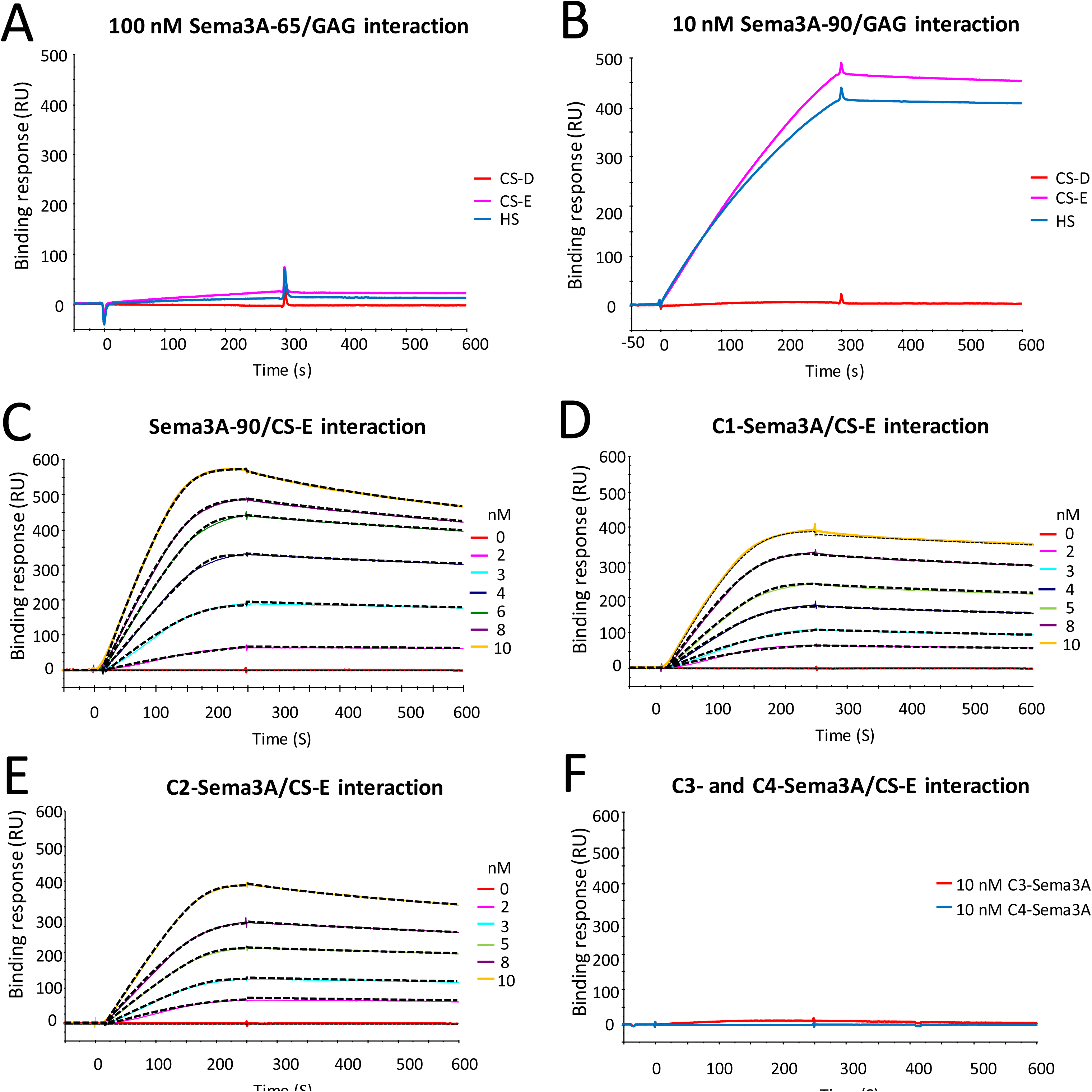
Sema3A binding to immobilized GAGs analysed by SPR. A: Overlay of sensorgrams showing binding of Sema3A-65 (100 nM) to immobilized CS-E (blue), CS-D (red) and HS (green). B: Overlay of sensorgrams showing binding of wild type Sema3A-90 (10 nM) to immobilized CS-E (pink), CS-D (red) and HS (blue). C-E: Wild type Sema3A-90 (C), C1-Sema3A (D) and C2-Sema3A binding to CS-E at 10, 8, 6, 5, 4, 3, 2 and 0 nM (color code as indicated). F: C3-Sema3A (red) and C4-Sema3A (blue) binding to CS-E at 10 nM. Proteins were injected for 4.2 min at 60 µl/min, after which running buffer alone was injected. Binding responses, in resonance units (RU), were recorded as a function of time. The association and dissociation phases across all Sema3A concentrations were fit to a 1:1 binding model including a mass transfer step, using the biaeval 3.1 software, to extract rate and equilibrium binding constants (see Table 1). Colored curves show the experimental data and dotted lines the results of the fit.

### Semaphorin 3A selectively interacts with CS-E and HS through its clusters 3 and 4

We next examined the binding of wild type and mutant Sema3A to different GAGs, including CS-D, CS-E and HS. To this end, we adopted a solid-phase assay and used SPR-based real-time monitoring to measure the binding of Sema3A to biotinylated GAGs immobilized in a streptavidin-displaying dextran matrix on sensor chips. As the biotin was conjugated at the reducing end of the GAG chains, GAG immobilization via streptavidin effectively mimics the matrix or cell membrane-anchored GAGs as, for example, in PNNs. Using this system, we first observed that Sema3A-65, at 100 nM, did not bind to any of the GAGs (Fig. 2A) while 10 nM of Sema3A-90 exhibited strong binding to both HS and CS-E, but not to CS-D (Fig. 2B).

It is common knowledge that the percentages of sulfated glycans in GAGs purified from natural sources are highly variable. To better define the GAGs used for the binding assay, we analyzed their disaccharide composition using RPIP-HPLC. Compositional analysis of CS-D and CS-E yielded remarkably different results (Supp mat Fig. 3). While the CS-E exhibited a clear majority (~70%) of disulfated E-units (Δdi-4S6S), CS-D was found to comprise a significant proportion of non-sulfated (~20%) and Δdi-6S monosulfated disaccharides (~50%), the disulfated D-unit (Δdi-2S6S) accounting for only ~30% of the polysaccharide composition. As CS chains do not feature S-domain like organization, such as that observed in vertebrate HS, it can be speculated, based on a stochastic distribution of the disaccharides along the polysaccharide chains, that occurrence of clusters of consecutive E-units on CS-E will be significantly greater than that of D-units on CS-D (see below). With more than two third of the CS-E disaccharide being disulfated, a homogeneous distribution would gave rise to an alternation of two disulfated disaccharides and one nonsulfated disaccharide along the chain (i.e. E-E-x-E-E-x), whereas in CS-D, each disulfated disaccharide (one third of the chain) would typically be spaced apart by two mono-or non-sulfated disaccharides (i.e. D-x-x-D-x-x). Therefore, we could not rule out the possibility that the absence of Sema3A binding to the CS-D polysaccharide observed by SPR could be due to the rarity of suitable binding sites on these chains, rather than a structural specificity of E (GlcA-GalNAc4S6S)-over D (GlcA2S-GalNAc6S)-units for the interaction with Sema3A.

**Figure 3:**
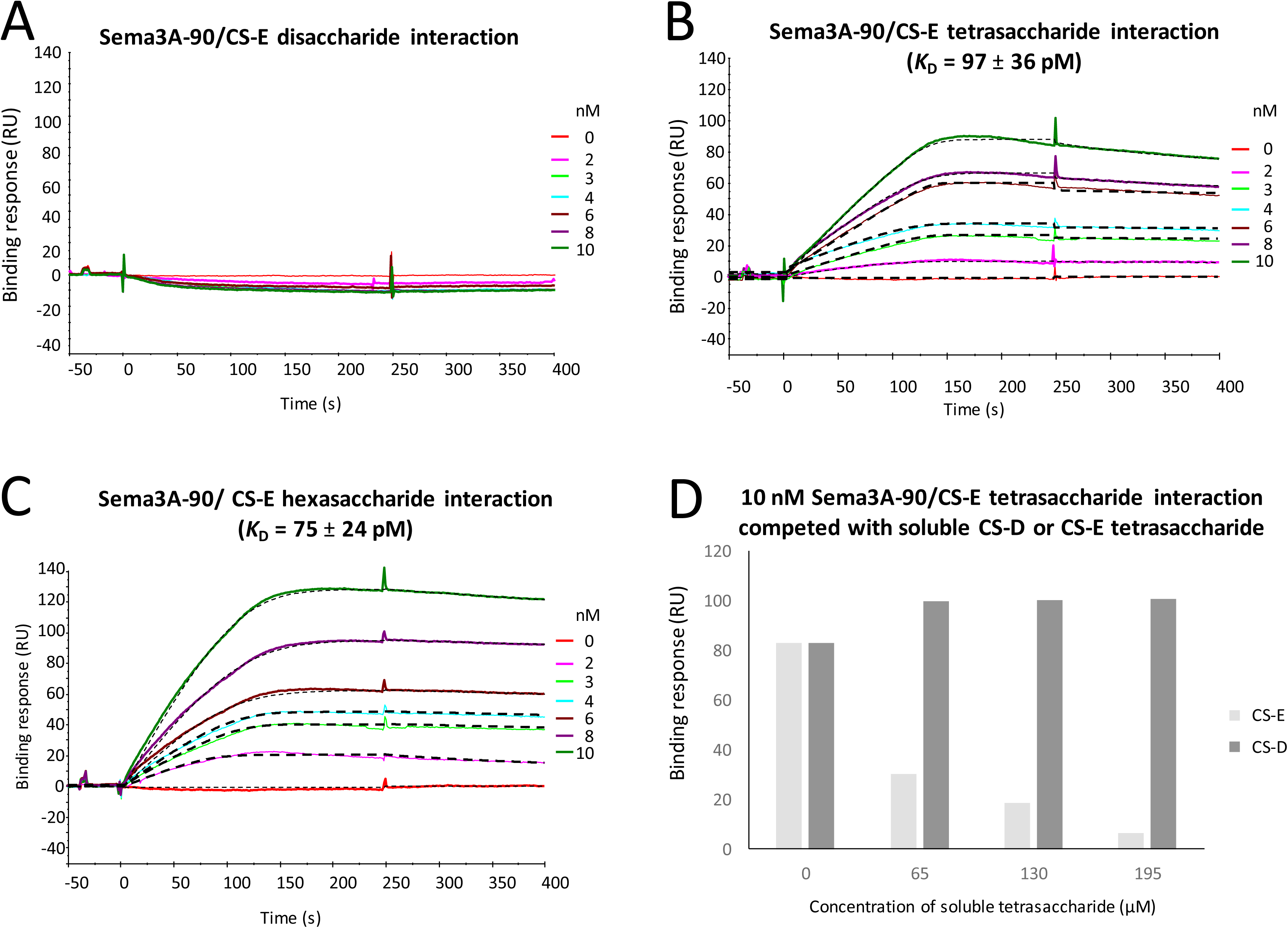
Sema3A binding to size-defined CS-E oligosaccharides. Sema3A (0 to 10 nM, as indicated) was injected over the sensor surface functionalized with synthetic biotinylated CS-E di-(A), tetra-(B) or hexa-(C) saccharides. The binding responses were recorded as described for Figure 2.D: Sema3A (10 nM) was preincubated with a range of concentrations of synthetic CS-E or CS-D tetrasaccharides and injected over a CS-E tetrasaccharide functionalized sensor. The binding responses at equilibrium (in RU) were plotted as a function of the concentration of CS-E (black) or CS-D (grey) tetrasaccharides in the solution-phase.

To further assess the contribution of the different basic clusters in CS-E binding, ranges of concentrations of wild type and mutant Sema3A were flowed across the CS-E surface and the resulting binding curves were fitted to a Langmuir binding model that included a mass transfer step (see Fig. 2C-F for data and fits, and Table 1 for results). These analyses demonstrated that WT-, C1-and C2-Sema3A featured very strong and similar binding (*K*_D_ < 0.1 nM) for CS-E, contributed both by a fast association rate constant and a low dissociation rate constant. Clusters 1 and 2 are thus not involved in the interaction. On the contrary, C3-and C4-Sema3A both demonstrated a complete absence of binding signal in the concentration range investigated, indicating the significance of clusters 3 and 4 in CS-E binding. A possible explanation is that clusters 3 and 4 each would bind independently yet rather weakly to CS-E, such that the presence of both clusters enhances binding substantially via multivalecy effect.

**Table 1:** Kinetic constants of Sema3A/CS-E interaction.

| | $k_{\text{on}}$ ($10^7 \text{ M}^{-1}\text{s}^{-1}$) | $k_{\text{off}}$ ($10^{-4} \text{ s}^{-1}$) | $K_D$ (pM) |
| --- | --- | --- | --- |
| WT Sema3A-90 | $1.1 \pm 0.3$ | $5.3 \pm 1.0$ | $47 \pm 2$ |
| C1-Sema3A | $1.5 \pm 0.6$ | $6.2 \pm 0.8$ | $41 \pm 3$ |
| C2-Sema3A | $6.9 \pm 0.2$ | $5.6 \pm 2.0$ | $82 \pm 8$ |

### A CS-E tetrasaccharide is the minimal motif that binds to Sema3A

To identify the minimal oligosaccharide length and the exact structure required for binding of Sema3A to CS-E, we used multi-step and stereo-controlled synthetic chemistry to generate a collection of size-and sulfation pattern-defined CS oligosaccharides. CS-E di-, tetra-, or hexa-saccharides were first chemically synthesized and biotinylated as previously described [33] and immobilized in a streptavidin-displaying dextran film on a SPR sensor chip. Conditions in this assay were chosen such that each streptavidin molecule harbors several copies of the oligosaccharide. Injection of Sema3A over these size-defined oligosaccharides demonstrated that Sema3A does not interact with disaccharides, but binds well to tetra-and hexa-saccharides of CS-E, with *K*_D_ values of 31 ± 9 and 36 ± 0.4 pM, respectively (Fig. 3A-C). The binding constants of Sema3A for the tetra-and hexa-saccharides are thus almost identical to those for the full length CS-E chains, indicating that a (GlcA-GalNAc4S6S)_2_ motif is likely representing the Sema3A binding epitope, a result consistent with the above described CS-E chain analysis.

The compositional analysis demonstrated a large variety of sulfated glycan composition in CSs purified from natural sources (Supp mat Fig. 3). In order to compare the Sema3A binding to disulfated CS-D and CS-E, we also chemically prepared non-biotinylated CS-D and CS-E tetrasaccharides [34] and used them in competition experiments. In this assay, Sema3A was pre-incubated with a range of concentrations (0 – 195 µM) of the tetrasaccharides CS-D or CS-E before injection over a sensor chip functionalized with CS-E tetrasaccharides. The result clearly showed that at the concentrations where CS-E tetrasaccharide inhibited over 90% of the binding, CS-D tetrasaccharide did not inhibit the Sema3A/CS-E binding (Fig. 3D). This finally demonstrates the specificity of the interaction between Sema3A and CS-E (i.e., its strict sulfation pattern dependence).

Interestingly, this competition assay also showed that µM concentrations of CS-E tetra-saccharides in the soluble phase are required to effectively compete for Sema3A binding to CS-E tetrasaccharides immobilized in the dextran matrix (Fig. 3D), whereas the direct binding experiments had shown effective binding of Sema3A to CS-E tetrasaccharides (Fig. 3B) and polysaccharides (Fig. 2C) in the dextran matrix at sub-nM concentrations. This striking difference indicates that binding to matrix-bound CS-E is enhanced by multivalent interactions. This interpretation is plausible considering that the dextran matrix presents multiple GAG chains in close proximity, and that Sema3A forms a disulfide-bonded dimer. It is tempting to suggest that such multivalency enables Sema3A to discriminate different forms of ECM according to their GAG presentation. For example, binding to PNNs may be favored over the loose ECM in the brain owing to different CS-E densities in these two matrices.

### Sema3A cross-links and rigidifies CS-E and HS matrices

We next investigated what could be the functional significance of the Sema3A/CS-E interaction. Previously, we reported that various HS binding signaling proteins rigidify and cross-link GAG chains and as such, could affect the supramolecular organization of HS in the extracellular space [35, 36]. The above-described evidence for multivalent interaction between Sema3A and CS-E tetrasaccharide clusters implies that Sema3A dimers are also able to engage simultaneously with more than one GAG chain, suggesting the protein is able to cross-link GAGs. We also noticed that expression of Sema3A in HEK293-6E cells strongly induced cell clustering (Sup mat Fig. 4), suggesting that Sema3A may simultaneously bind to and cross-link multiple GAG polysaccharides on the cell surface and thus agglutinate cells via their GAG coats.

**Figure 4:**
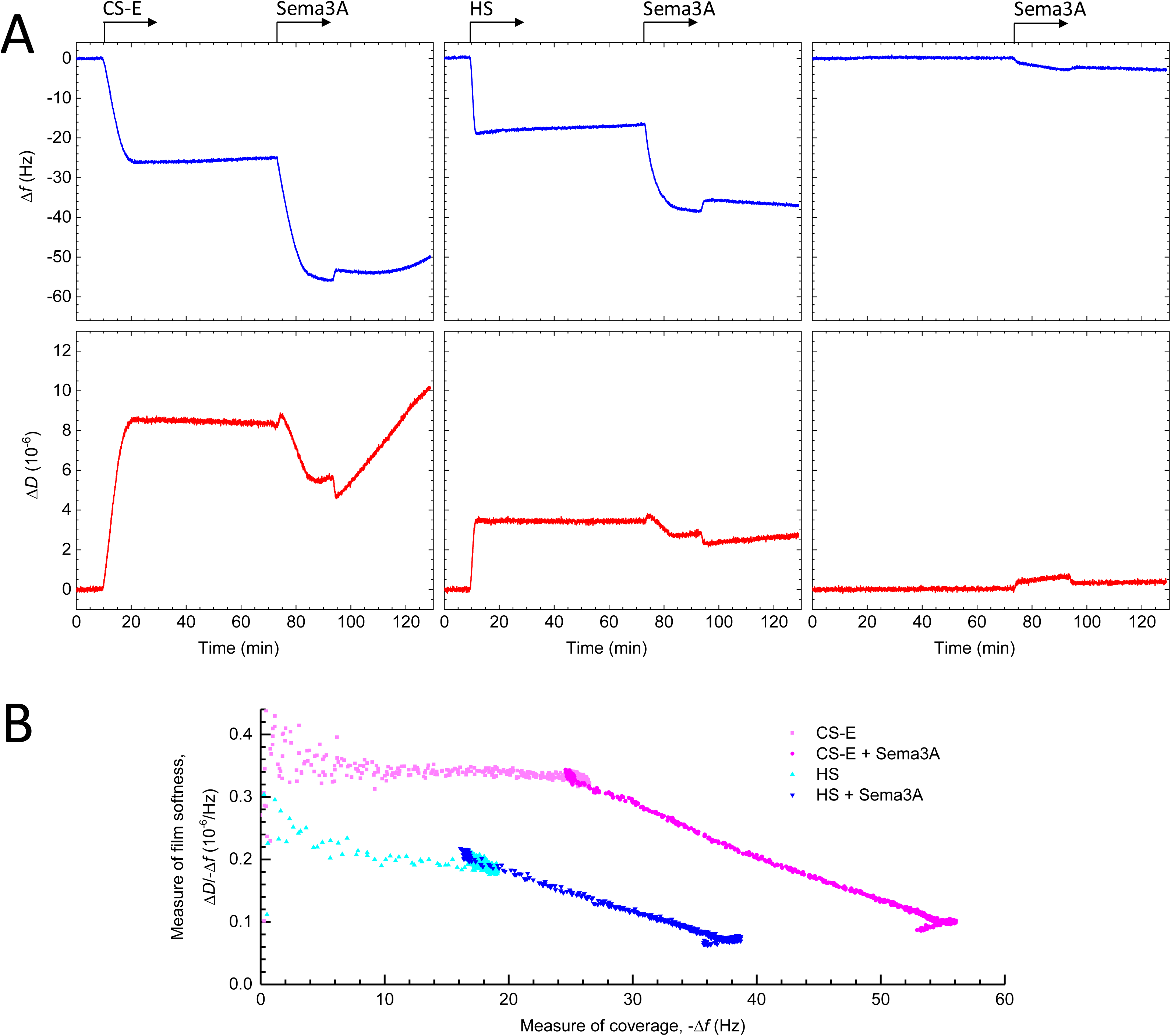
Sema3A rigidifies CS-E and HS films. A-C: QCM-D data (top: frequency shift, Δ*f*; bottom: dissipation shift, Δ*D*) for the formation of films of CS-E (A) and HS (B) or without any GAGs (negative control; C), and the subsequent binding of Sema3A-90 at 20 μg/ml. The GAG films were formed on a dense monolayer of streptavidin on a silica-supported lipid bilayer; arrows on top of the graph indicate the start and duration of incubation with GAGs and Sema3A; throughout all remaining times the sensor surface was exposed to plain working buffer. Note that the minor yet rapid (i.e. within 2 min) changes seen in Δ*f* and (more prominently) Δ*D* upon the exchanges from plain working buffer to Sema3A, and back, do not reflect any changes on the surface but result from an increase in the viscosity and/or density of the surrounding solution owing to the presence of added salt in the Sema3A solution. D: Parametric plot of Δ*D*/-Δ*f* versus -Δ*f* for Seam3A-90 binding to CS-E and HS (as indicated; data from A and B). Δ*D*/-Δ*f* is a measure of the elastic compliance (or ‘softness’) of GAG films, and -Δ*f* is a measure of GAG and Sema3A surface coverage; the plot demonstrates the substantial film rigidification (or decrease in softness) upon Sema3A binding.

To further test this possibility, we analysed by QCM-D the effect of Sema3A binding on GAG film softness. In this assay. GAG chains were anchored via their reducing ends to the QCM-D sensor surface, akin to their anchorage in proteoglycans, and the mean distance between anchor points was chosen sufficiently short such that adjacent GAG chains could readily encounter each other. Sema3A-90, but not Sema3A-65, bound to CS-E and HS as evidenced by the clear decrease in resonance frequency (Δ*f*) upon exposure of the protein to the GAG films (Fig. 4A; Sup mat Fig. 5), consistent with the earlier SPR data. Notably, Sema3A binding induced a strong decrease in dissipation (Δ*D*) (Fig. 4A) indicating that Sema3A substantially rigidifies CS-E and HS films. The parametric plot (Fig. 4B) shows the Δ*D*/-Δ*f* ratio, which is a measure of the elastic compliance (or ‘softness’) of the GAG films, against -Δ*f* as a measure of GAG and Sema3A surface coverage. This plot illustrates that whilst the bare CS-E film is softer than the bare HS film (as expected, owing to the CS-E chains being longer than the HS chains thus likely forming thicker yet less dense films) the softness decreases (i.e. rigidity increases) to a similar level for both films upon binding of Sema3A. A plausible explanation for the observed rigidification effect is that Sema3A cross-links GAGs chains, in line with the SPR and cell agglutination data.

**Figure 5:**
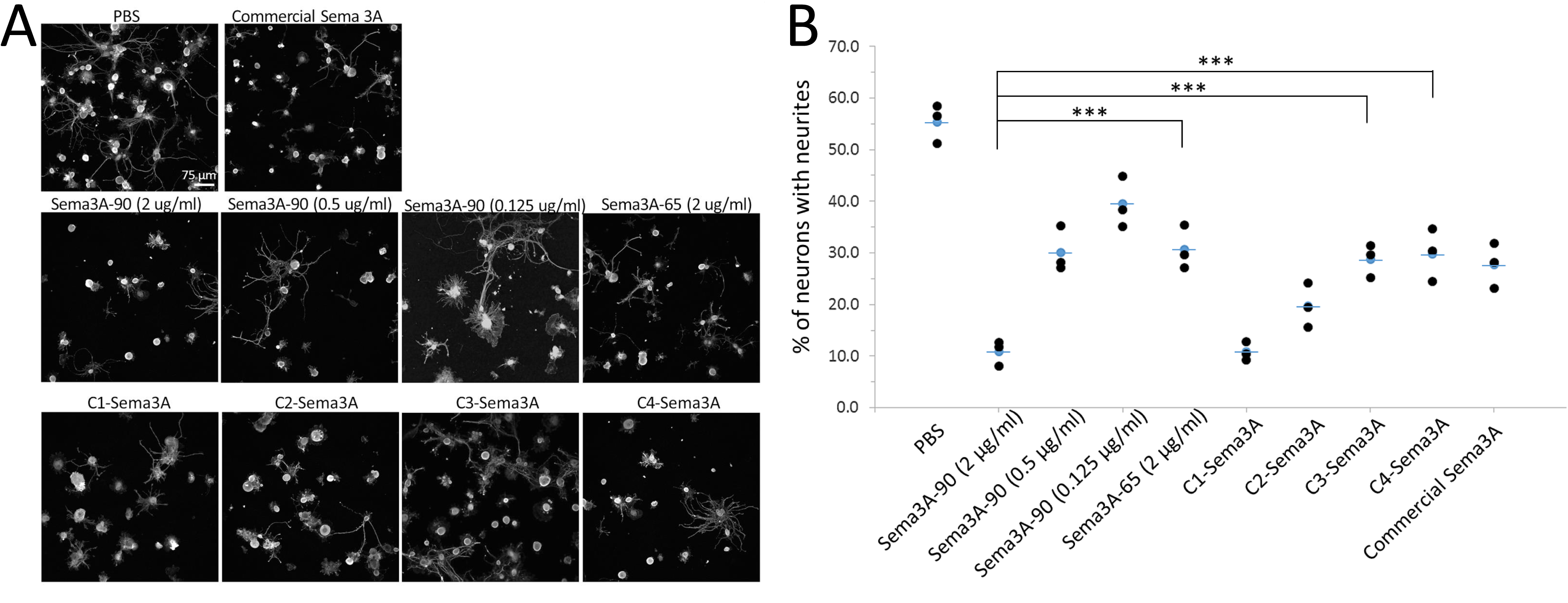
The C-terminal domain of Sema3A is important for the full inhibitory effect of Sema3A on neurons. A: Adult DRG neurons were cultured in the presence of different Sema3A isoforms and mutants (as indicated), and stained for βIII tubulin to reveal the neurites. B: A graph showing the quantification of percentage of neurones bearing neurites in the presence of the indicated Sema3A proteins.

### CS binding sites are involved in Sema3A-induced inhibition of neurite outgrowth

We have previously shown that Sema3A binding potentiates the inhibitory properties of CSs to DRG neurons. Here, we assessed the effect of Sema3A and its mutants on the inhibition of neurite outgrowth on DRG neurons, to check the activity of purified Sema3A and to assess the importance of identified GAG-binding sequences in Sema3A signalling *in vitro*. Dissociated adult rat DRG neurons, which lose their neurites during their extraction from parent tissues, were cultivated in the presence of 100 ng/ml of NGF. Neurons were then treated with PBS (negative control), 2 µg/ml of commercial Sema3A (positive control; a construct with a stabilizing mutation to prevent furin cleavage at site 1; see Fig. 1), different concentrations of wild type Sema3A-90, or 2 µg/ml of Sema3A-65, C1-Sema3A, C2-Sema3A, C3-Sema3A or C4-Sema3A (Fig. 5A). The number of neurons projecting neurites in each condition was quantified and presented as percentage of neurons with neurites (Fig. 5B). The results show firstly that wild type Sema3A-90 inhibits neurite outgrowth, thus confirming that the purified protein is active. Secondly, this inhibition effect is Sema3A concentration-dependent (Fig. 5B). Sema3A elicits significant inhibitory effects at 2 µg/ml. This concentration was thus used to analyse the effect of other proteins. Results showed that Sema3A-90 was more inhibitory (three times) than Sema3A-65, indicating the importance of the C-terminal domains to potentiate the inhibitory effect of Sema3A. C1-and C2-Sema3A induced the same level of inhibition as the wild type Sema3A-90, while C3-and C4-Sema3A displayed the same level of inhibition as Sema3A-65 (Fig. 5B). These data suggest that the C-terminal domain of Sema3A, and in particular the CS binding site, is important for the full inhibitory effect of Sema3A on neurons.

## DISCUSSION

We have successfully expressed and purified full-length Sema3A for the first time, without any stabilizing mutations. Using full-length Sema3A along with Sema3A-65 and basic cluster mutants, we have shown that the C-terminal domain of Sema3A is responsible for binding to GAGs, including CS-E and HS. The interaction requires a tetrasaccharide as the minimal saccharide size, shows very high avidity in the context of GAG-rich matrices, induces crosslinking of GAG chains, and rigidifies GAG films. We further identified the basic clusters 3 and 4 at the C-terminal as responsible for CS-E binding.

Sema3A features two conserved basic cleavage sites for furin-like pro-protein convertases, located within the PSI domain and in the C-terminal basic tail, respectively. Cleavage of the former site, which releases the 65 kDa Sema domain substantially diminishes the chemorepulsive property of Sema3A [22]. It was previously shown that overexpressed Sema3A is also proteolytically processed at the first internal furin cleavage site (RTRR, residues 552-555 in the PSI domain) and that the C-terminal was removed during the purification, unless mutations (R554A and R555A) were introduced within the furin site [9]. When using HEK293 cells as a cellular system to produce recombinant Sema3A we observed, consistent with the above-mentioned studies, that most of the secreted protein present in the conditioned medium displayed a molecular mass of 65 kDa, corresponding to the furin processed and less active form. However, we found that the majority of the secreted protein remained associated to the cell surface, from which it could be displaced using high salt concentrations. The cell-surface bound protein displayed the molecular mass (90 kDa) expected from the full-length protein and we made use of this observation to prepare large amounts of intact Sema3A, enabling us to investigate the structure/function relationships of the Sema3A C-terminal domain, in the context of the full length, wild type protein. I should be noted however that we could not totally excluded that our preparation did not contained a small proportion of Sema 3A cleaved at the second furin cleavage site. Such a cleavage would indeed reduce the molecular mass of 1.4 kDa only (see Fig. 1). In addition, the intact and cleaved proteins can assemble into homo-dimers as well as heterodimers, thus producing 3 distinct dimer forms.

Class-3 semaphorins are the only vertebrate semaphorins that are expressed as secreted proteins, while other vertebrate semaphorins are transmembrane or membrane anchored proteins, which can occasionally be further processed into soluble forms by proteolytic cleavage. The observation that the cell surface triggers an almost quantitative accumulation of full length Sema3A, which thus exists essentially in a bound form, is intriguing. It suggests that even Sema3A, as a member of the soluble class-3 semaphorin family, requires some form of immobilization for maximal activity. In contrast to semaphorins from the other subfamilies, however, Sema3A is not confined to the cell membrane but may be immobilized anywhere in the extracellular space through (multivalent) GAG binding.

While a full length Sema3A is essential for the binding to Nrp1 and thus triggers growth cone collapse, the function of cleaved Sema3A is less clear. With the use of adult rat DRG neurons, our results show that strong neurite inhibition is only observed with wild type Sema3A-90, C1-Sema3A and C2-Sema3A, while Sema3A-65, C3-Sema3A and C4-Sema3A all demonstrate weaker inhibition. This suggests the C-terminal domain of Sema-3A is important for its full function. This agrees with a previous observation using embryonic chick DRG neurons where neurite projections were inhibited with the full length Sema3A and weaker with the C-terminal deletion mutant [22]. A recent paper by Serini et al. [37] has shown that, outside the CNS, a C-terminal deletion mutant of Sema3A induces endothelial cell migration similar to wild type Sema3A through direct interaction with plexinA4. Whether this mechanism also applies to the function of Sema3A in the CNS remains to be elucidated.

Our results suggest that the retention of full length Sema3A on the cell surface is dependent on the presence of GAGs such as CS-E or HS. In CNS, the majority of the sulfated GAGs belongs to CS, with mono-sulfated CSs as the prevalent forms [38]. The proportion of CS-E, when compared to other CS isoforms, gradually increases during CNS development and reaches ~1.4% upon birth [38, 39]. In PNNs, this proportion is further increased to beyond 2%. Although the presence of CS-E is low, it is crucial for the selective binding of some molecules. Prochiantz et al. [40] have previously suggested that CS-E in PNNs allows for the specific binding of orthodenticle homeobox protein 2 (Otx2) and thus controls the maturation of parvalbumin neurons in the visual cortex. In collaboration with Verhaagen et al., we have also shown Sema3A is enriched on the surface of PNN positive neurons [12, 13]. Here, our results further characterize the Sema3A-CS-E binding, and the high avidity of the interaction suggests that the presence of 2% CS-E will be sufficient to lead to Sema3A enrichment on the neuronal surface.

Having the full length and wild type Sema3A in hand, we used two independent experimental approaches, GAG-protein cross-linking followed by N-terminal sequencing and mutagenesis coupled to SPR binding analysis, to demonstrate that, of the four basic clusters in the C-terminal domain of Sema3A, only clusters 3 and 4 contribute to GAG binding. Kinetic analysis of the binding data first showed that Sema3A associates to CS-E with very high avidity (*K*_D_ in the sub nM range), contributed by both a high association rate constant and a low dissociation rate constant. The very strong stability of the resulting complex can be explained by the dimeric nature of Sema3A, which thus features a total of four GAG-binding clusters (i.e. two per monomer) for multivalent binding to GAG-rich matrices.

While Sema3A-90 demonstrated a very strong binding CS-E or HS using SPR, it is important to note the need of µM concentrations of CS-E tetrasaccharides to compete for Sema3A binding to immobilized CS-E. This implies that the multivalency of Sema3A allows and facilitates simultaneous binding to multiple GAG chains presented on surfaces, including those in our solid phase assays and in PNNs. The exact arrangement of the four clusters upon GAG binding remains unknown, but we here propose several possibilities (schematically shown in Fig. 6) that would all be consistent with the available data. Of particular note is the close proximity of the disulfide bond to the two clusters 3, which implies a close proximity of at least two GAG-binding clusters in the Sema3A dimer. This spatial constraint could well explain the potency of Sema3A to bring GAG chains together, as required for efficient cross-linking and rigidification of GAG matrices.

**Figure 6:**
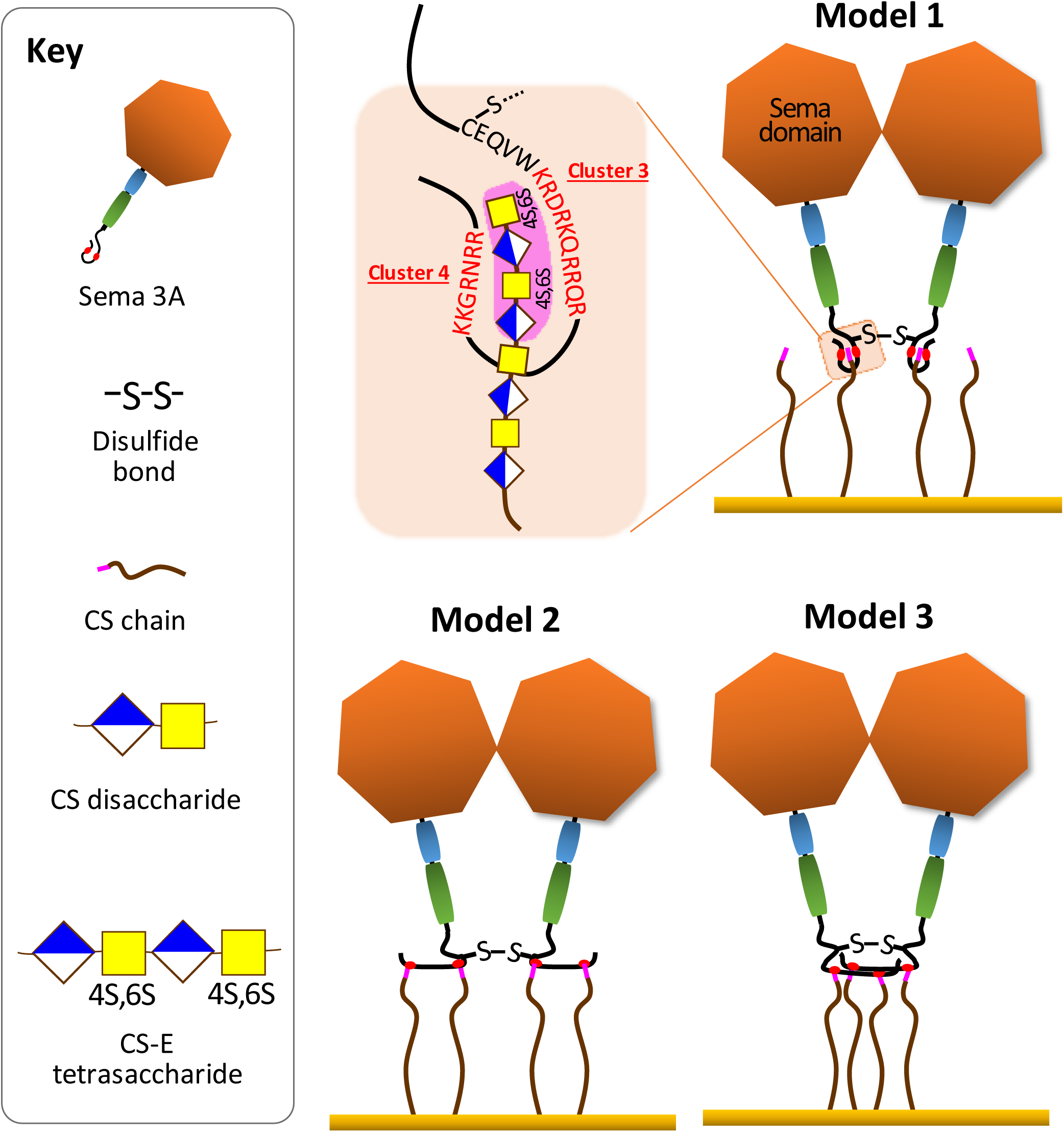
Molecular mechanisms of multivalent GAG-binding and GAG cross-linking. The schematics show putative mechanisms consistent with all experimental data in the context of the Sema3A dimer. Note the spatial proximity of the C723-C723 disulfide bond (-S-S-) to the GAG-binding clusters 3.

Interestingly, cluster 3 (KRDRKQRRQR) is very similar to a basic sequence (RKQRRER) found in Otx2, a protein which is also involved in PNN formation and plasticity. This basic sequence mediates accumulation of Otx2 into PNNs, through binding to CS [40] raising the possibility that Sema3A and Otx2 could compete with one another, an aspect which merits further investigation as it could help to better understand the interplay of these two proteins in PNN assembly, dynamics and function.

GAG binding cluster 4 is also one of the two furin cleavage sites of Sema3A. Such dual function has been observed already, for example in Sindbis virus, where furin protease cleavage of the recognition sequence mediates virion attachment to cell surface HS [41]. In HIV, within the domain spanning the gp120/gp41 junction, a GAG binding and furin recognition sequence also overlap [42]. While a peptide encompassing this region is only a poor furin substrate in the absence of heparin, addition of the polysaccharide importantly increases the cleavage suggesting a functional relationship between GAG binding and furin activity [43]. It would thus be interesting to investigate whether the C-terminal furin recognition sequence in Sema3A, cleavage of which enhances the biological activity of the protein [22], is also better processed upon binding to CS-E.

Examining the GAG structural requirements for Sema3A recognition, we showed that Sema3A binds to CS-E but not to CS-D. These results are consistent with previous reports obtained using polysaccharides isolated from squid and shark cartilage respectively [12]. These materials, however, have not been precisely characterized so that the analysis of the binding data remains uncertain with regards to their exact structure. Compositional analysis of CS-D and CS-E indeed indicated that beyond the difference in the nature of their representative disaccharide units (GlcA2S-GalNac6S versus GlcA-GalNac4S6S) they feature highly distinct overall charge, with only 30 % of disulfated disaccharides for CS-D versus 70 % for CS-E. In addition, the exact distribution of the disaccharides along the GAG chains is unknown. To better understand the structural basis of the CS-E and D difference in Sema3A binding capacity, we thus turned our effort toward synthetic chemistry and generated oligosaccharides that are defined in their size and sulfation pattern. This approach enabled us to demonstrate that a CS-E (GalNac-GlcA4S6S)_2_ tetrasaccharide motif represents the specific Sema3A binding epitope, whereas a CS-D motif of the same length (GlcA2S-GalNac6S)_2_ is totally devoid of binding activity. The observation that Sema3A binds to a CS-E motif as short as a tetrasaccharide with an affinity almost identical to that of a full-length CS-E chain opens the possibility of using such small ligand to interfere with the interaction *in vivo*. In that perspective, it would be interesting to combine two tetrasaccharides through an appropriate linker, fitting the Sema3A dimer geometry, a strategy successfully developed for interferon-γ, another homodimeric cytokine [44, 45].

## ACKNOWLEDGEMENTS

We would like to thank Prof. Joost Verhaagen (the Netherlands Institute of Neurosciences) for constructive discussions, Jean-Claude Jacquinet (the Institut de Chimie Organique et Analytique) for help with the chondroitin sulfate chemical synthesis, Jean-Pierre Andrieu (the Institut de Biologie Structurale) for the N-terminal sequencing and Lewis Adams (University of Leeds) for support with pilot QCM-D experiments.

## ABBREVIATIONS

Sema: Semaphorin
GAG: Glycosaminoglycan
CS: Chondroitine sulfate
HS: Heparan sulfate
PNN: Perineuronal nets
ECM: Extracellular Matrix

## Reference

1. Worzfeld T & Offermanns S (2014) Semaphorins and plexins as therapeutic targets. Nature Reviews Drug Discovery 13:603.

2. Yazdani U & Terman JR (2006) The semaphorins. Genome biology 7(3):211.

3. Goodman CS, Kolodkin AL, Luo Y, Püschel W, & Raper JA (1999) Unified nomenclature for the semaphorins/collapsins. Semaphorin Nomenclature Committee. Cell 97(5):551–552.

4. Antipenko A, Himanen JP, van Leyen K, Nardi-Dei V, Lesniak J, Barton WA, Rajashankar KR, Lu M, Hoemme C, Puschel AW, & Nikolov DB (2003) Structure of the semaphorin-3A receptor binding module. Neuron 39(4):589–598.

5. Janssen BJ, Robinson RA, Perez-Branguli F, Bell CH, Mitchell KJ, Siebold C, & Jones EY (2010) Structural basis of semaphorin-plexin signalling. Nature 467(7319):1118–1122.

6. Siebold C & Jones EY (2013) Structural insights into semaphorins and their receptors. Seminars in cell & developmental biology 24(3):139–145.

7. Nogi T, Yasui N, Mihara E, Matsunaga Y, Noda M, Yamashita N, Toyofuku T, Uchiyama S, Goshima Y, Kumanogoh A, & Takagi J (2010) Structural basis for semaphorin signalling through the plexin receptor. Nature 467(7319):1123–1127.

8. Wang Y, He H, Srivastava N, Vikarunnessa S, Chen YB, Jiang J, Cowan CW, & Zhang X (2012) Plexins are GTPase-activating proteins for Rap and are activated by induced dimerization. Science signaling 5(207):ra6.

9. Janssen BJC, Malinauskas T, Weir GA, Cader MZ, Siebold C, & Jones EY (2012) Neuropilins lock secreted semaphorins onto plexins in a ternary signaling complex. Nature Structural & Molecular Biology 19:1293.

10. Zimmer G, Schanuel SM, Burger S, Weth F, Steinecke A, Bolz J, & Lent R (2010) Chondroitin sulfate acts in concert with semaphorin 3A to guide tangential migration of cortical interneurons in the ventral telencephalon. Cerebral cortex (New York, N.Y.: 1991) 20(10):2411–2422.

11. De Wit J, De Winter F, Klooster J, & Verhaagen J (2005) Semaphorin 3A displays a punctate distribution on the surface of neuronal cells and interacts with proteoglycans in the extracellular matrix. Molecular and cellular neurosciences 29(1):40–55.

12. Dick G, Tan CL, Alves JN, Ehlert EM, Miller GM, Hsieh-Wilson LC, Sugahara K, Oosterhof A, van Kuppevelt TH, Verhaagen J, Fawcett JW, & Kwok JC (2013) Semaphorin 3A binds to the perineuronal nets via chondroitin sulfate type E motifs in rodent brains. The Journal of biological chemistry 288(38):27384–27395.

13. Vo T, Carulli D, Ehlert EM, Kwok JC, Dick G, Mecollari V, Moloney EB, Neufeld G, de Winter F, Fawcett JW, & Verhaagen J (2013) The chemorepulsive axon guidance protein semaphorin3A is a constituent of perineuronal nets in the adult rodent brain. Molecular and cellular neurosciences 56:186–200.

14. Testa D, Prochiantz A, & Di Nardo AA (2019) Perineuronal nets in brain physiology and disease. Seminars in cell & developmental biology 89:125–135.

15. Romberg C, Yang S, Melani R, Andrews MR, Horner AE, Spillantini MG, Bussey TJ, Fawcett JW, Pizzorusso T, & Saksida LM (2013) Depletion of perineuronal nets enhances recognition memory and long-term depression in the perirhinal cortex. The Journal of neuroscience: the official journal of the Society for Neuroscience 33(16):7057–7065.

16. Yang S, Cacquevel M, Saksida LM, Bussey TJ, Schneider BL, Aebischer P, Melani R, Pizzorusso T, Fawcett JW, & Spillantini MG (2015) Perineuronal net digestion with chondroitinase restores memory in mice with tau pathology. Experimental neurology 265:48–58.

17. Pizzorusso T, Medini P, Berardi N, Chierzi S, Fawcett JW, & Maffei L (2002) Reactivation of ocular dominance plasticity in the adult visual cortex. Science (New York, N.Y.) 298(5596):1248–1251.

18. Boggio EM, Ehlert EM, Lupori L, Moloney EB, De Winter F, Vander Kooi CW, Baroncelli L, Mecollari V, Blits B, Fawcett JW, Verhaagen J, & Pizzorusso T (2019) Inhibition of Semaphorin3A Promotes Ocular Dominance Plasticity in the Adult Rat Visual Cortex. Molecular neurobiology. doi: 10.1007/s12035-019-1499-0. [Epub ahead of print].

19. Ma CW, Kwan PY, Wu KL, Shum DK, & Chan YS (2019) Regulatory roles of perineuronal nets and semaphorin 3A in the postnatal maturation of the central vestibular circuitry for graviceptive reflex. Brain structure & function 224(2):613–626.

20. Bosch KD, Bradbury EJ, Verhaagen J, Fawcett JW, & McMahon SB (2012) Chondroitinase ABC promotes plasticity of spinal reflexes following peripheral nerve injury. Experimental neurology 238(1):64–78.

21. Corredor M, Bonet R, Moure A, Domingo C, Bujons J, Alfonso I, Perez Y, & Messeguer A (2016) Cationic Peptides and Peptidomimetics Bind Glycosaminoglycans as Potential Sema3A Pathway Inhibitors. Biophysical journal 110(6):1291–1303.

22. Adams RH, Lohrum M, Klostermann A, Betz H, & Puschel AW (1997) The chemorepulsive activity of secreted semaphorins is regulated by furin-dependent proteolytic processing. The EMBO journal 16(20):6077–6086.

23. Klostermann A, Lohrum M, Adams RH, & Puschel AW (1998) The chemorepulsive activity of the axonal guidance signal semaphorin D requires dimerization. The Journal of biological chemistry 273(13):7326–7331.

24. Adi SD, Eiza N, Bejar J, Shefer H, Toledano S, Kessler O, Neufeld G, Toubi E, & Vadasz Z (2019) Semaphorin 3A Is Effective in Reducing Both Inflammation and Angiogenesis in a Mouse Model of Bronchial Asthma. Frontiers in immunology 10:550.

25. Lavi N, Kessler O, Ziv K, Nir-Zvi I, Mumblat Y, Eiza N, Paran Y, Brenner B, Vadasz Z, & Neufeld G (2018) Semaphorin-3A inhibits multiple myeloma progression in a mouse model. Carcinogenesis 39(10):1283–1291.

26. L’Abbe D, Bisson L, Gervais C, Grazzini E, & Durocher Y (2018) Transient Gene Expression in Suspension HEK293-EBNA1 Cells. Methods in molecular biology (Clifton, N.J.) 1850:1–16.

27. Vives RR, Crublet E, Andrieu JP, Gagnon J, Rousselle P, & Lortat-Jacob H (2004) A novel strategy for defining critical amino acid residues involved in protein/glycosaminoglycan interactions. The Journal of biological chemistry 279(52):54327–54333.

28. Henriet E, Jager S, Tran C, Bastien P, Michelet JF, Minondo AM, Formanek F, Dalko-Csiba M, Lortat-Jacob H, Breton L, & Vives RR (2017) A jasmonic acid derivative improves skin healing and induces changes in proteoglycan expression and glycosaminoglycan structure. Biochimica et biophysica acta. General subjects 1861(9):2250–2260.

29. Saesen E, Sarrazin S, Laguri C, Sadir R, Maurin D, Thomas A, Imberty A, & Lortat-Jacob H (2013) Insights into the mechanism by which interferon-gamma basic amino acid clusters mediate protein binding to heparan sulfate. Journal of the American Chemical Society 135(25):9384–9390.

30. Thakar D, Migliorini E, Coche-Guerente L, Sadir R, Lortat-Jacob H, Boturyn D, Renaudet O, Labbe P, & Richter RP (2014) A quartz crystal microbalance method to study the terminal functionalization of glycosaminoglycans. Chemical communications (Cambridge, England) 50(96):15148–15151.

31. Lopin C & Jacquinet JC (2006) From polymer to size-defined oligomers: an expeditious route for the preparation of chondroitin oligosaccharides. Angewandte Chemie International ed. in English 45(16):2574–2578.

32. Migliorini E, Thakar D, Sadir R, Pleiner T, Baleux F, Lortat-Jacob H, Coche-Guerente L, & Richter RP (2014) Well-defined biomimetic surfaces to characterize glycosaminoglycan-mediated interactions on the molecular, supramolecular and cellular levels. Biomaterials 35(32):8903–8915.

33. Jacquinet JC & Lopin-Bon C (2015) Stereocontrolled preparation of biotinylated chondroitin sulfate E di-,tetra-,and hexasaccharide conjugates. Carbohydrate research 402:35–43.

34. Jacquinet JC, Lopin-Bon C, & Vibert A (2009) From polymer to size-defined oligomers: a highly divergent and stereocontrolled construction of chondroitin sulfate A, C, D, E, K, L, and M oligomers from a single precursor: part 2. Chemistry (Weinheim an der Bergstrasse, Germany) 15(37):9579–9595.

35. Dyer DP, Migliorini E, Salanga CL, Thakar D, Handel TM, & Richter RP (2017) Differential structural remodelling of heparan sulfate by chemokines: the role of chemokine oligomerization. Open biology 7(1).

36. Migliorini E, Thakar D, Kuhnle J, Sadir R, Dyer DP, Li Y, Sun C, Volkman BF, Handel TM, Coche-Guerente L, Fernig DG, Lortat-Jacob H, & Richter RP (2015) Cytokines and growth factors cross-link heparan sulfate. Open biology 5(8).

37. Gioelli N, Maione F, Camillo C, Ghitti M, Valdembri D, Morello N, Darche M, Zentilin L, Cagnoni G, Qiu Y, Giacca M, Giustetto M, Paques M, Cascone I, Musco G, Tamagnone L, Giraudo E, & Serini G (2018) A rationally designed NRP1-independent superagonist SEMA3A mutant is an effective anticancer agent. Science translational medicine 10(442).

38. Deepa SS, Carulli D, Galtrey C, Rhodes K, Fukuda J, Mikami T, Sugahara K, & Fawcett JW (2006) Composition of perineuronal net extracellular matrix in rat brain: a different disaccharide composition for the net-associated proteoglycans. The Journal of biological chemistry 281(26):17789–17800.

39. Kitagawa H, Tsutsumi K, Tone Y, & Sugahara K (1997) Developmental regulation of the sulfation profile of chondroitin sulfate chains in the chicken embryo brain. The Journal of biological chemistry 272(50):31377–31381.

40. Beurdeley M, Spatazza J, Lee HH, Sugiyama S, Bernard C, Di Nardo AA, Hensch TK, & Prochiantz A (2012) Otx2 binding to perineuronal nets persistently regulates plasticity in the mature visual cortex. The Journal of neuroscience : the official journal of the Society for Neuroscience 32(27):9429–9437.

41. Klimstra WB, Heidner HW, & Johnston RE (1999) The furin protease cleavage recognition sequence of Sindbis virus PE2 can mediate virion attachment to cell surface heparan sulfate. Journal of virology 73(8):6299–6306.

42. Crublet E, Andrieu JP, Vives RR, & Lortat-Jacob H (2008) The HIV-1 envelope glycoprotein gp120 features four heparan sulfate binding domains, including the co-receptor binding site. The Journal of biological chemistry 283(22):15193–15200.

43. Pasquato A, Dettin M, Basak A, Gambaretto R, Tonin L, Seidah NG, & Di Bello C (2007) Heparin enhances the furin cleavage of HIV-1 gp160 peptides. FEBS letters 581(30):5807–5813.

44. Lubineau A, Lortat-Jacob H, Gavard O, Sarrazin S, & Bonnaffe D (2004) Synthesis of tailor-made glycoconjugate mimetics of heparan sulfate that bind IFN-gamma in the nanomolar range. Chemistry (Weinheim an der Bergstrasse, Germany) 10(17):4265–4282.

45. Sarrazin S, Bonnaffe D, Lubineau A, & Lortat-Jacob H (2005) Heparan sulfate mimicry: a synthetic glycoconjugate that recognizes the heparin binding domain of interferon-gamma inhibits the cytokine activity. The Journal of biological chemistry 280(45):37558–37564.

